# Cancer exosomes induce tumor neo-neurogenesis potentiating tumor growth

**DOI:** 10.1101/247452

**Authors:** Marianna Madeo, Paul L. Colbert, Daniel W. Vermeer, Christopher T. Lucido, Elisabeth G. Vichaya, Aaron J. Grossberg, Jacob T. Cain, DesiRae Muirhead, Alex P. Rickel, Zhongkui Hong, William C. Spanos, John H. Lee, Robert Dantzer, Paola D. Vermeer

**Affiliations:** Cancer and Immunotherapies Group, Sanford Research, 2301 East 60^th^ St north, Sioux Falls, SD 57104.; Pediatrics and Rare Disease Group, Sanford Research, 2301 East 60^th^ St north, Sioux Falls, SD 57104.; Department of Symptom Research, MD Anderson Cancer Center, 1515 Holcombe Blvd, Unit 384, Houston, TX 77030.; Department of Radiation Oncology, MD Anderson Cancer Center, 1400 Pressler St, Unit 1422, Houston, TX 77030.; Department of Pathology, LCM Pathologists, Sanford Health, 1305 West 18^th^ St, Sioux Falls, SD 57105.; Biomedical Engineering Program, University of South Dakota, 4800 North Career Ave, Sioux Falls, SD 57107.; Sanford Ears, Nose and Throat, 1310 West 22^nd^ St, Sioux Falls, SD 57105.; NantKwest, 9920 Jefferson Blvd, Culver City, CA 90232.

## Abstract

Patients with densely innervated tumors do worse than those with less innervated cancers. We hypothesize that neural elements are acquired by a tumor-induced process, called neo-neurogenesis. Here, we use PC12 cells in a simple system to test this hypothesis. PC12 cells extend processes, called neurites, only when appropriately stimulated. Using this system, we show that patient tumors release vesicles (exosomes) which induce PC12 neurite outgrowth. Using a cancer mouse model, we show that tumor cells compromised in exosome release grow slower and are less innervated than controls indicating a contribution of innervation to disease progression. We find that neo-neurogenesis is mediated in part by the axonal guidance molecule, EphrinB1, contained in exosomes. These findings support testing EphrinB1 blockers to inhibit tumor innervation and improve survival.

**One Sentence Summary:** Tumors release exosomes which not only promote their own innervation but also potentiate their growth.

## Main Text

### Introduction

Innervated tumors are more aggressive than less innervated ones (*1–7*). For instance, in prostate cancer, recruitment of nerve fibers to cancer tissue is associated with higher tumor proliferative indices and a higher risk of recurrence and metastasis (*5*). Denervation studies in pre-clinical and genetically engineered mouse cancer models support a functional contribution of neural elements in disease progression (*8*, *9*) (*10*). These studies strongly indicate that the nervous system is not a bystander but instead an active participant in carcinogenesis and cancer progression. However, a mechanistic understanding of how tumors obtain their neural elements remains unclear. Tumors may acquire innervation by growing within innervated tissues; in other words, neural elements are already present within the microenvironment and the tumor acquires them by default. However, the clinical findings that some tumors of the same tissue are more innervated than others indicate instead an active, tumor-initiated process, similar to neo-angiogenesis and lymphangiogenesis. The possibility that tumors invoke their own innervation, termed neo-neurogenesis, has not been extensively explored (*11*, *12*).

Extracellular release of neurotrophic factors [e.g. nerve growth factor (NGF)] by tumor cells can contribute to cancer progression (*13*, *14*). While such a direct mechanism likely contributes to tumor innervation, tumors release additional components which may directly promote neo-neurogenesis. Among these are extracellular vesicles such as exosomes. Exosomes are 30-150nm vesicles that package a rich cargo (proteins, DNA, RNA, lipids). Because they are generated by invagination of endocytic vesicles, surface cargo protein topology is preserved as is, presumably, biological activity. Exosomes are released into the extracellular milieu by most, if not all, cells (*15*) and function as vehicles of intercellular communication (*16*) (*17*). Mounting evidence supports the hypothesis that cancer cells utilize exosomes to induce/promote metastasis and tumor tolerance (*17*) (*18*). Here, we show that tumor released exosomes mediate neo-neurogenesis in cancer which contributes to disease progression.

We utilize a murine model of human papillomavirus induced (HPV+) oropharyngeal squamous cell carcinoma (OPSCC) which consists of C57Bl/6 oropharyngeal epithelial cells stably expressing HPV16 viral oncogenes, E6 and E7, H-Ras and luciferase (mEERL cells) (*19*) (*20-22*). The HPV16 E6 oncoprotein interacts with the cellular phosphatase and tumor suppressor, PTPN13; this interaction results in PTPN13’s degradation (*20*, *21*). This is relevant because PTPN13 interacts with many cellular proteins including EphrinB1 which is also a phosphatase substrate (*23*). EphrinB1 is a single pass transmembrane protein ligand that binds and activates the Eph receptor tyrosine kinases. Furthermore, EphrinB1 itself becomes phosphorylated and initiates its own downstream signaling (*24*). In HPV-infected cells, PTPN13 expression is compromised and thus EphrinB1 phosphorylation persists and contributes to an aggressive disease phenotype (*25*) (*26*). During development, EphrinB1 functions as an axonal guidance molecule (*27*) (*28*). Here we show that tumor released exosomes package EphrinB1 and stimulate neurite outgrowth of PC12 cells *in vitro*. Compromise of EphrinB1 expression or function significantly attenuates this activity. Moreover, exosomes purified from human squamous cell carcinoma cell lines and from head and neck cancer patient plasma and tumor also package EphrinB1 as exosomal cargo and harbor neurite outgrowth activity. Consistent with these *in vitro* findings, mEERL tumors over-expressing EphrinB1 are significantly more innervated than tumors with compromised EphrinB1 function or expression. In addition, mEERL tumors genetically compromised in exosome release are sparsely innervated *in vivo* and grow slower than controls. Taken together, these data indicate that tumor released exosomes contribute to neo-neurogenesis and that exosomal EphrinB1 potentiates this activity.

### Results

#### Patient exosomes induce neurite outgrowth

We tested whether patient HNSCCs are innervated by immunohistochemically (IHC) staining formalin-fixed paraffin embedded tumor tissue for β-III tubulin, a neuron specific tubulin isoform. β-III tubulin positive fibers are found coursing throughout the tissue indicating these tumors are indeed innervated (Figure 1A, “Nerve twigs”). These β-III tubulin positive nerve “twigs” cannot be confused with perineural invasion (PNI). (*29*, *30*). PNI refers to tumor invading into nerves along the perineural space; neo-neurogenesis refers to nerves invading into tumor. Within the perineural sheath, β-III tubulin positive fibers are packed tightly together in a very organized manner (Figure 1A, “nerve bundle”). The β-III tubulin positive nerve fibers we have identified are instead coursing as individual, unorganized twigs lacking a perineural sheath (Figure 1A “nerve twigs”). Additional IHC staining shows that HNSCCs are negative for tyrosine hydroxylase (sympathetic marker) and VIP (parasympathetic marker) but positive for TRPV1 (sensory marker) (Figure 1A) indicating sensory neo-innervation of tumor.

**Fig. 1.**
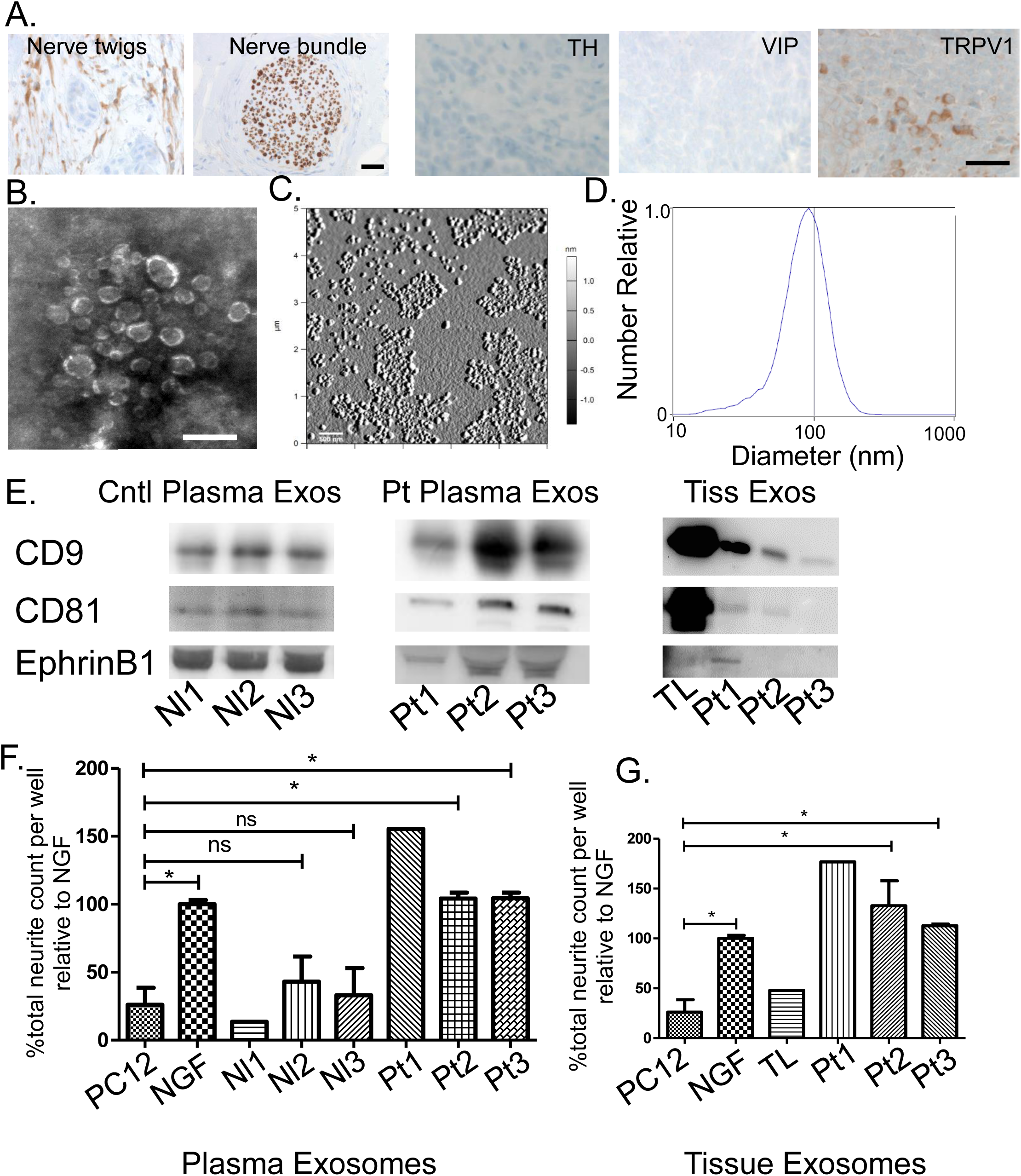
Head and neck cancer patient derived exosomes induce neurite outgrowth. A) HNSCC: nerve “twigs” and “bundle” IHC for β-III tubulin (scale bar, 50µm), Tyrosine Hydroxylase (TH), Vasoactive Intestinal Polypeptide (VIP) and Transient Receptor Potential Vanilloid-type one (TRPV1)(scale bar, 20µm). Brown, immunostain; blue, counterstain. Exosome scanning electron micrograph (scale bar, 200nm) (B), atomic force microscopy amplitude trace (scale bar, 500nm) (C) and nanoparticle tracking analysis (D). E) Control (Cntl), patient (Pt) plasma and tissue (Tiss) exosomes (Exos) western blots. Neurite outgrowth following plasma (F) or tissue (G) exosome stimulation. Nl1, Nl2, Nl3 controls; Pt1, Pt2, Pt3, patients; TL, tonsil. Student’s t-test. *, p<0.05; **, p<0.001. Error bars, standard deviation.

Prior to testing the contribution of exosomes to neo-neurogenesis, we performed validations of our differential ultracentrifugation exosome purification technique (*31*). For human blood samples, exosomes were purified from plasma. For human tissue samples, exosomes were similarly purified from conditioned media collected after 48 hours in culture. Scanning electron microscopic analysis of our exosome preparations purified from normal donor plasma yielded vesicles consistent in shape and size (30-150nm) with exosomes (Figure 1B). Additionally, atomic force microscopy confirmed a 65-110 nm size (Figure 1C) and nanoparticle tracking analysis for counting and sizing exosomes also indicated a size distribution consistent with exosomes (Figure 1D) (*32*). Taken together, these data indicate that our purification method yields vesicles consistent in size and shape with exosomes.

To test our hypothesis that tumor released exosomes induce neo-neurogenesis, we utilized PC12 cells, a rat pheochromocytoma cell line, as an *in vitro* screen. When stimulated with NGF (100 ng/ml), PC12 cells differentiate into neuron-like cells and extend neurites (*33*). We collected 10ml of blood along with matched tumor tissue from three head and neck cancer patients (patient samples Pt1, Pt2, Pt3). We similarly collected blood from 3 healthy volunteers (Nl1, Nl2, Nl3) as well as adult tonsil tissue (TL). The tonsil was chosen as control tissue since the majority of HPV+ OPSCCs arise in the tonsil. Exosomes were purified, quantified by BCA protein assay and further validated by western blot analysis for the exosome markers CD9 and CD81 (Figure 1E). To test whether they harbor neurite outgrowth activity, PC12 cells were treated with 3 g exosomes, fixed 48 hours later and immunostained for β-III tubulin. Neurite outgrowth was quantified using the CellInSight CX7 High Content Analysis Platform and the number of neurites compared. The exosome yield from patient Pt1 was low allowing for analysis of only one replicate while quantities from Pt2 and Pt3 were sufficient for technical replicates. Consistent with the literature, we found that untreated PC12 cells extend very few β-III tubulin positive neurites while those stimulated with NGF do so robustly. Exosomes from all three patients (both plasma and tumor) stimulated significant neurite outgrowth of PC12 cells while exosomes from normal plasma and tonsil had minimal neurite outgrowth activity (Figure 1 F, G). These data indicate that exosomes from head and neck cancer patients harbor neurite outgrowth activity that is absent in healthy controls.

#### mEERL exosomes induce neurite outgrowth

To model the process of neo-neurogenesis, mice were injected with mEERL cells into the hind limb; tumors were later harvested at endpoint, fixed, embedded and IHC stained for β-III tubulin, TH, VIP and TRPV1. Similar to patient HNSCCs, mEERL tumors harbored β-III tubulin positive nerve twigs that were sensory in nature (TRPV1 positive) (Figure 2A). To test whether mEERL released exosomes contribute to neo-neurogenesis, cells were cultured *in vitro*, exosomes purified from conditioned media and tested on PC12 cells. To test the function of EphrinB1 in this process, we generated EphrinB1 modified mEERL cell lines. Stable over-expression of wild-type EphrinB1 has been previously characterized and is referred to as mEERL EphrinB1 (*26*). Utilizing CRISPR/Cas9, we genetically engineered mEERL cell lines compromised in EphrinB1 function or expression. EphrinB1 deleted cells are denoted as mEERL EphrinB1 Null1 or Null2 while extracellularly deleted EphrinB1 cells are denoted as mEERL EphrinB1ΔECD. The characterization of these mEERL EphrinB1 CRISPR lines is presented in Supplemental Figures 1 and 2. Exosomes from mEERL parental cells significantly induced neurite outgrowth of PC12 cells. Over-expression of EphrinB1 increased this activity. Interestingly, exosomes from mEERL EphrinB1ΔECD, Null 1 and Null 2 cells retain the ability to induce neurite outgrowth (Figure 2B). Taken together, these data indicate that mEERL released exosomes promote neurite outgrowth and that while EphrinB1 is not required for this activity, it significantly potentiates it.

**Fig. 2.**
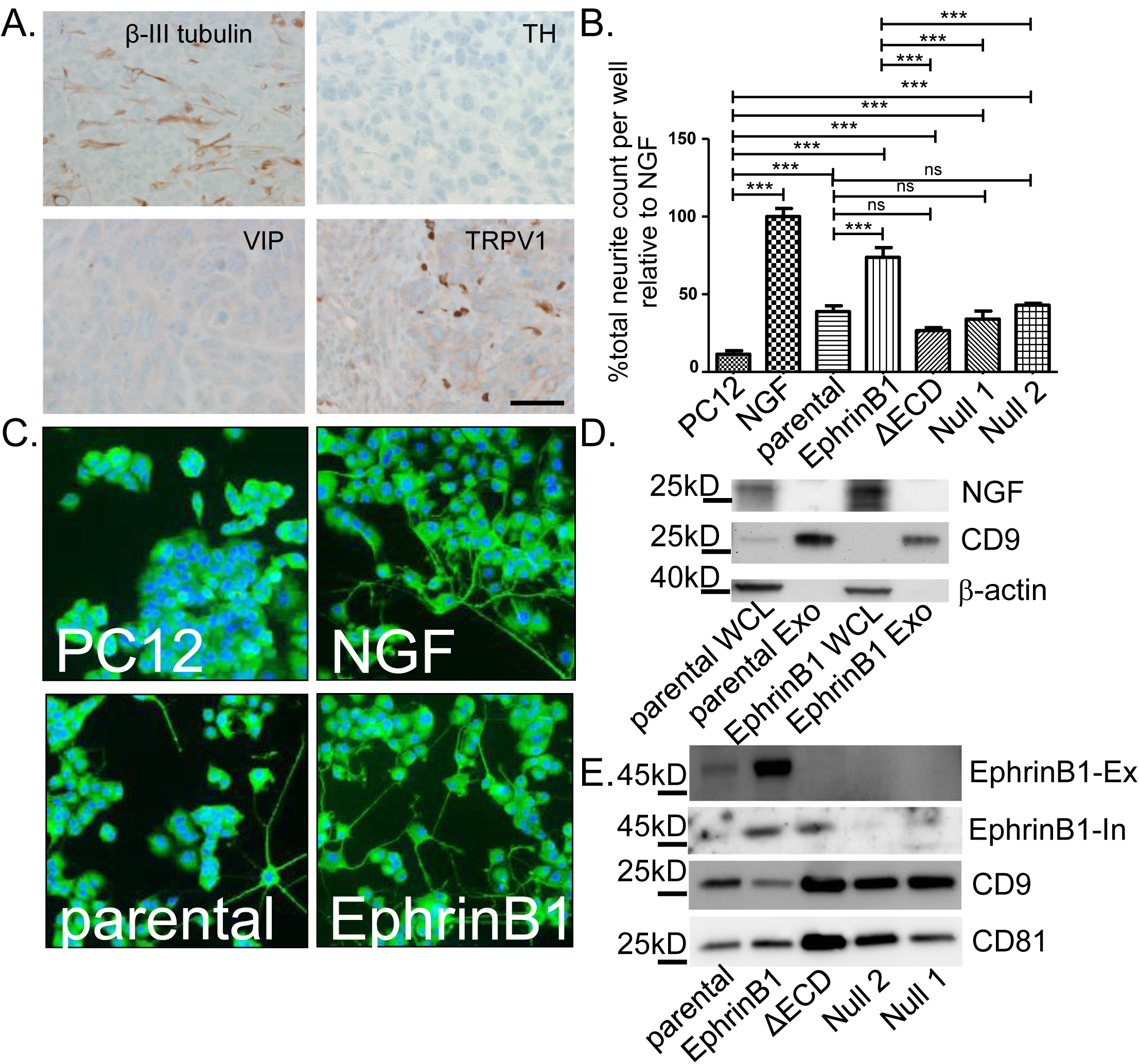
mEERL exosomes induce neurite outgrowth. A) IHC (brown) of mEERL tumor. Tyrosine Hydroxylase (TH), Vasoactive Intestinal Polypeptide (VIP) and Transient Receptor Potential Vanilloid-type one (TRPV1). Scale bar, 20µm. B) Neurite outgrowth quantification following exosome treatment. Statistical analysis: one-way ANOVA, post hoc analysis by Tukey test. ***, p<0.05; ns, not significant. N=4 replicates/condition; experiment repeated twice. C) β-III tubulin positive immunofluorescent (green) PC12 cells following exosome stimulation. Blue, DaPi nuclear stain. D) Western blot analysis. Whole cell lysate (WCL); Exosomes (Exo). E) Western blot analysis of exosomes. EphrinB1-Ex, EphrinB1 extracellular epitope antibody. EphrinB1-In, EphrinB1 intracellular epitope antibody. Error bars, standard deviation.

#### Exosomes induce neurite outgrowth without NGF

As mEERL cells can produce NGF (*34*), we analyzed whole cell lysates and purified exosomes from mEERL parental and EphrinB1 cells by western blot for NGF. We confirmed that mEERL parental and EphrinB1 over-expressing cells produce NGF (present in whole cell lysate, WCL), but showed that it is not packaged within CD9+ exosomes (Figure 2D). These data indicate that NGF is not required for exosome-mediated neurite outgrowth activity. Given that exosomes purified from mEERL EphrinB1 cells potentiate neurite outgrowth of PC12 cells, we tested whether it was packaged as exosome cargo. Western blot analysis of exosomes indicated that EphrinB1 is indeed packaged within exosomes (Figure 2E) which is consistent with the published literature (*35*) (Exocarta.org). Moreover, while the extracellular domain of EphrinB1 is absent in mEERL EphrinB1ECD exosomes, the intracellular domain remains as cargo. Importantly, EphrinB1 was also found in patient exosomes (Figure 1E). As with our human exosome validation, we similarly validated exosomes purified from mEERL cells and found them to be likewise consistent in size and shape with exosomes (see Supplemental Figure 3A-C).

**Fig. 3.**
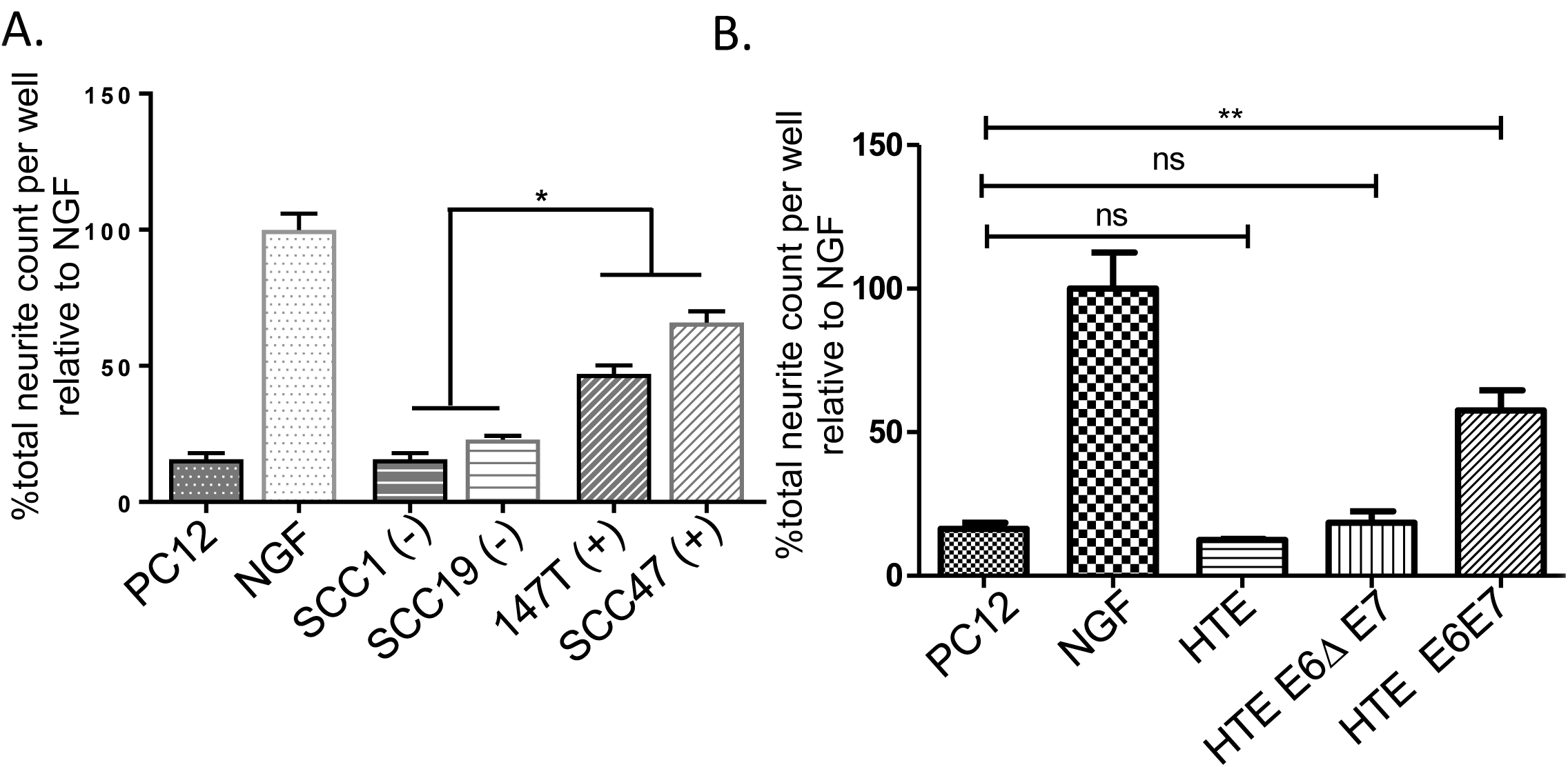
HPV status contributes to neurite outgrowth activity. Neurite outgrowth following exosome stimulation with: A) HPV negative (−) or positive (+) human cells; statistical analysis by one-way ANOVA comparing the four lines with post hoc Tukey test for differences between HPV- and + groups. *, p<0.001. B) HTE, human tonsil epithelia; HTE E6ΔE7, cells expressing HPV16 E6 deleted of its PDZBM(Δ) and E7; HTE E6E7, cells expressing HPV16 E6 and E7. Statistical analysis by student’s t-test. **, p<0.05; ns, not significant. Error bars, standard deviation. All assays: N=4 replicates/condition; experiment repeated twice.

Recent published studies suggest that more stringent methods are critical for eliminating other vesicles and cellular debris from exosome purifications (*36*). One such method requires the addition of density gradient centrifugation (*37*). To test whether this more stringent methodology purifies exosomes with neurite outgrowth activity, conditioned media from mEERL EphrinB1 cells was collected, subjected to differential ultracentrifugation and subsequently to density gradient centrifugation. Fifteen fractions were collected and fractions 4-13 were analyzed by western blot for CD9 and CD81. Consistent with the published literature, we CD9+ and CD81+ vesicles were present in fraction 8 (*38*, *39*); Exosomes purified by differential ultracentrifugation alone (“crude” sample) were also CD9+/CD81+ (Supplemental Figure 3D). Next, to determine which fractions retained neurite outgrowth activity, fractions 4, 5, 8 and 13 were tested on PC12 cells. “Crude” exosomes were also tested. While fraction 8 and “crude” exosomes demonstrate neurite outgrowth activity, CD9 negative fractions 4, 5 and 13 lacked this activity (Supplemental Figure 3E). These data indicate that inclusion of density gradient centrifugation concentrates CD9+/CD81+exosomes to a single fraction which retains neurite outgrowth activity. The data also indicate that neurite outgrowth activity is retained in EphrinB1 positive CD9+/CD81+ exosomes.

#### High-Risk HPV E6 and neurite outgrowth

Thus far, we have tested neurite outgrowth activity from mEERL cells or their derivatives, all of which are HPV+. We next tested exosomes from two HPV+ (SCC47 and 93-VU- 147T-UP-6) and two HPV- (SCC1 and SCC19) human squamous cell carcinoma cell lines on PC12 cells and found that the HPV- exosomes harbor significantly less neurite outgrowth activity than the HPV+ exosomes (Figure 3A). Since HPV16 induces OPSCC, it is considered a high risk HPV (*40*). Low risk HPVs rarely cause cancer. One important difference between high and low risk HPVs is found in their E6 proteins. Only high risk E6 contains a C-terminal PDZ binding motif (PDZBM) which significantly contributes to oncogenic transformation (*41*) (*42*). To test the contribution of HPV16 E6 to neurite outgrowth, we tested exosomes from primary human tonsil epithelia (HTE), as well as those stably expressing HPV16 E6 and E7 (HTE E6E7) or exosomes from cells in which the PDZBM of E6 has been deleted (HTE E6ΔE7). We found that expression of full length E6 together with E7 was sufficient to induce neurite outgrowth activity while deletion of E6’s PDZBM abrogated this effect (Figure 3B). Taken together, these data suggest that HPV16 E6 contributes to neurite outgrowth activity.

#### Exosomes promote tumor innervation and growth

To test whether EphrinB1 expression affects tumor innervation *in vivo*, mice were implanted with either mEERL parental, EphrinB1, ΔECD, Null1 or Null 2 cells. Ten days post-implantation, when tumors were palpable, they were harvested and whole tumor lysate subjected to western blot analysis for β-III tubulin. β-III tubulin signals were normalized to GAPDH and quantified by densitometry. EphrinB1 and EphrinB1ΔECD tumors harbor significantly more β-III tubulin compared to mEERL parental tumors (Figure 4A, 4B). This *in vivo* capacity to induce tumor innervation was different from *in vitro* where EphrinB1ΔECD exosomes induce significantly less neurite outgrowth from PC12 cells than mEERL parental exosomes (Figure 2B). This discrepancy likely reflects components within the tumor microenvironment (absent *in vitro*) that also affect tumor innervation. Similar to the *in vitro* data, Null 1 and Null 2 tumors were not different from mEERL parental (Figures 2B, 4A, 4B). Taken together, these data indicate that full length and truncated EphrinB1 are sufficient to potentiate tumor innervation *in vivo* while its complete deletion cannot.

**Fig 4.**
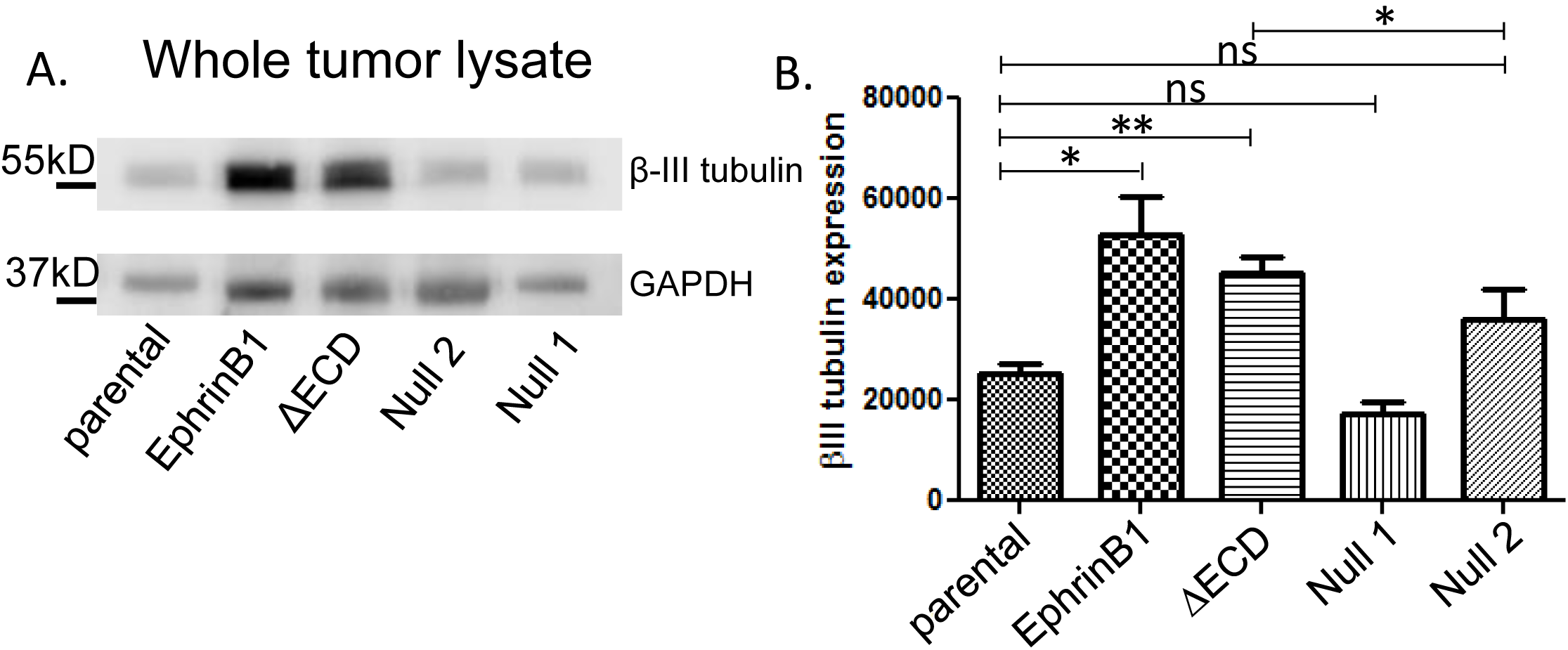
Mouse tumors are innervated *in vivo*. A) Western blot analysis for β-III tubulin and GAPDH for the indicated tumors. B) Densitometric quantification of β-III tubulin western blots in A. β-III tubulin signal was normalized to GAPDH. N= 4 tumors/condition were analyzed. Statistical analysis by student’s t-test; **, p<0.05; *, p<0.01; ns, not significant. Error bars, standard deviation.

To more stringently test the hypothesis that tumor released exosomes induce tumor innervation *in vivo* and to define its contribution to tumor growth, we utilized CRISPR/Cas9 to genetically modify Rab27A and/or Rab27B in mEERL parental cells. These two small GTPases contribute to exosome release and their knock-down compromises release of CD9+ exosomes (*43-45*). The clone generated is heterozygous for Rab27A and homozygous deleted for Rab27B (mEERL Rab27A^−/+^ Rab27B^−/−^). For characterization of this clone, see Supplemental Figure 4; exosome samples were normalized to producing cell number. Exosomes from mEERL parental and Rab27A^−/+^ Rab27B^−/−^ cells were purified and analyzed for CD9 and CD81 by western blot. Exosomes purified from the Rab27A^−/+^ Rab27B^−/−^ expressed less CD9 which is consistent with previous studies demonstrating a decreased capacity to release exosomes by cells compromised in Rab27A/B expression which is reflected by decreased CD9 expression (Figure 5A) (45). Nanoparticle tracking analysis confirmed the decreased capacity of mEERL Rab27A^−/+^ Rab27B^−/−^ cells to release exosomes (Figure 5B). Moreover, we labeled exosomes with CFDA-SE, a cell permeant fluorescein tracer, and quantified fluorescence. Exosomes from mEERL Rab27A^−/+^ Rab27B^−/−^ cells had decreased fluorescence relative to mEERL parental exosomes (Figure 5C). As a whole, these data confirm that modulation of Rab27A/B expression resulted in decreased exosome release. To test whether compromised exosome release affects neurite outgrowth of PC12 cells, mEERL parental and mEERL Rab27A^−/+^ Rab27B^−/−^ exosomes were normalized to producer cell number and equivalent volumes applied to PC12 cells. The neurite outgrowth activity of mEERL Rab27A^−/+^ Rab27B^−/−^ exosomes was significantly attenuated compared to that of exosomes from mEERL parental cells (Figure 5D). To test whether compromised exosome release alters innervation *in vivo*, mice were implanted with mEERL parental or Rab27A^−/+^ Rab27B^−/−^ cells and tumor growth monitored. Rab27A^−/+^ Rab27B^−/−^ tumors grew significantly slower than mEERL parental tumors *in vivo* (Figure 5E). Importantly, *in vitro* proliferation assays show that cell doubling time was not significantly different between mEERL parental and Rab27A^−/+^ Rab27B^−/−^ cells (Figure 5F). To determine whether this decreased capacity to release exosomes affects tumor innervation *in vivo*, tumors were harvested from mice and 30 g of whole tumor lysate quantified by western blot for β-III tubulin (Figure 5G). Due to the delayed growth of the mEERL Rab27A^−/+^ Rab27B^−/−^, tumors were harvested at 21 days. Consistent with our hypothesis, Rab27A^−/+^ Rab27B^−/−^ tumors were significantly decreased in β-III tubulin as compared to mEERL parental tumors (Figure 5H). These data support our hypothesis that tumor released exosomes contribute to neo-neurogenesis and also suggest that neo-neurogenesis affects tumor growth.

**Fig 5.**
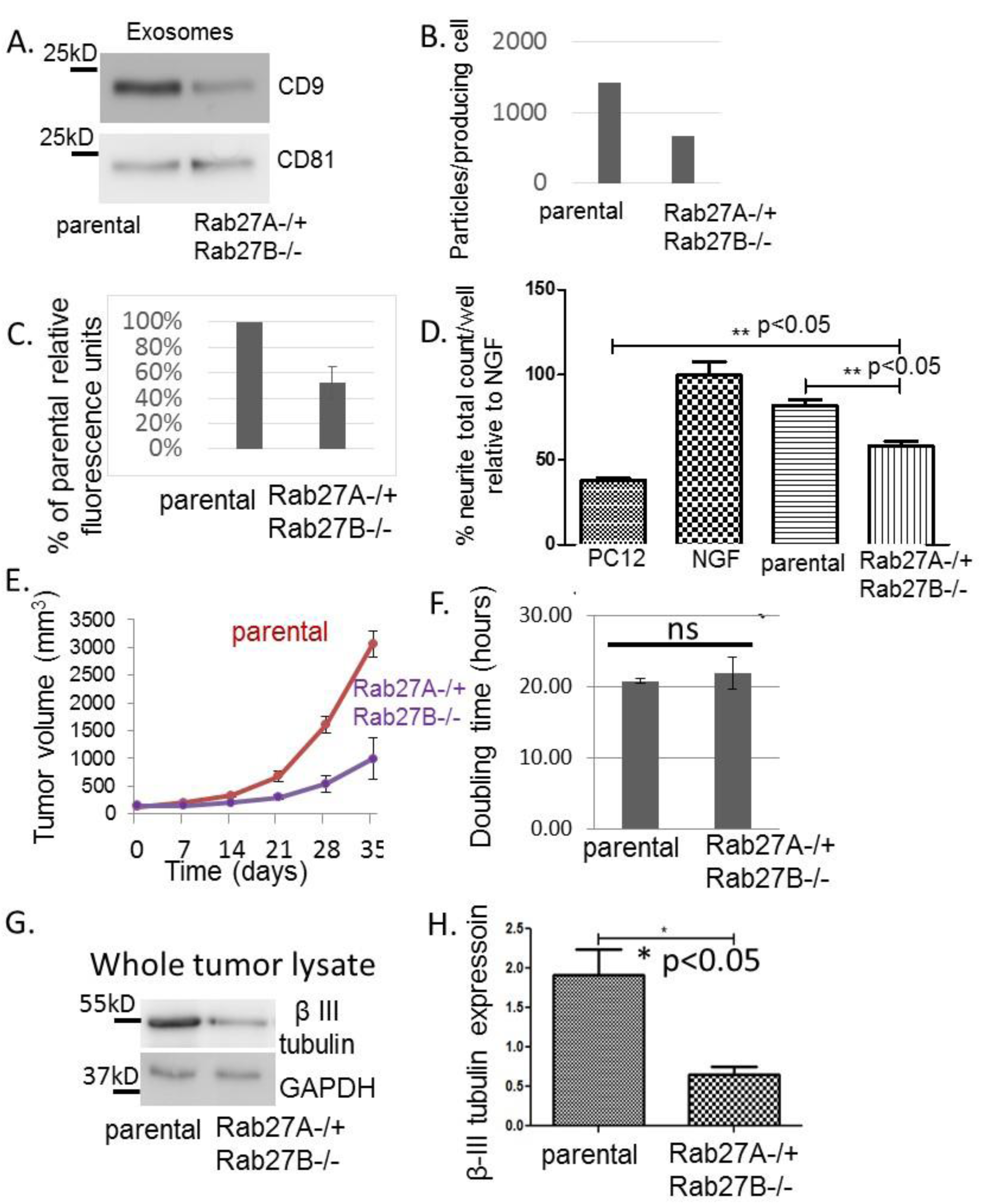
mEERL tumors compromised in exosome release are sparsely innervated and grow slower. A) Western blot of exosomes; repeated N=4 with similar results. B) Particles per producing cell number; repeated N=6 with similar results. C) Relative fluorescence units of CFDA-SE labeled exosomes. N=2 samples/condition. D) Neurite outgrowth following exosome stimulation. N=4 replicates/condition; experiment repeated twice. E) Tumor growth curves; N=7 mice/condition. F) Proliferation assay. Repeated N=3 times with similar results. G) Whole tumor lysate western blot. H) Densitometric quantification of G. Exosomes normalized to producing cell number. Statistical analysis by student’s t-test; ns=not significant; p values indicated; error bars, SEM.

#### Interpretations

Our findings propose a new mechanism for tumor-induced neo-neurogenesis. We show that human and mouse HPV+ HNSCCs are innervated *de novo* by TRPV1 positive sensory nerves. Moreover, while mEERL tumors secrete NGF, it is not required for neurite outgrowth activity in our *in vitro* assay nor is it packaged within exosomes. Mechanistically, packaging of full length EphrinB1 as exosome cargo significantly potentiates neurite outgrowth *in vitro* and tumor innervation *in vivo* and its deletion significantly attenuates both of these activities. Compromising release of CD9+ exosomes results in significantly decreased tumor growth and innervation *in vivo*. These pre-clinical studies are supported by findings with human HNSCC samples where HNSCC patient plasma and tumor exosomes harbor neurite outgrowth activity. Taken together, these data indicate that CD9+ exosomes released by HPV+ tumor cells promote tumor innervation and tumor growth *in vivo*. Exosomes containing EphrinB1 further potentiate this activity. HPV infection could modulate exosome cargo and, in this way, affect neurite outgrowth activity. Alternatively, the effects of HPV and EphrinB1 could be related. In HPV infected cells, E6’s interaction with PTPN13 results in the degradation of this phosphatase. As a consequence, EphrinB1 phosphorylation persists. Phosphorylated EphrinB1 interacts with binding partners which could then shuttle along with it into exosomes, a theory supported by our EphrinB1ΔECD data. If HPV’s contribution to neo-neurogenesis and disease progression is via this mechanism, it stands to reason that head and neck cancers would harbor either mutations in PTPN13 or EphrinB1 but not both. In fact, The Cancer Genome Atlas shows that PTPN13 and EphrinB1 alterations are mutually exclusive in HNSCC. This mutual exclusivity extends to breast, ovarian, prostate, liver, lung cancers. Thus, our findings could be significant for other cancers. The data also indicate other exosomal cargo including DNA, RNA, miRNA and lipids should be examined for their neurogenic activity. Whatever the combination of factors involved, it is clear that interventions targeting tumor exosome release or blocking the ability of nerves to respond to exosomes may be of therapeutic value. While this concept requires rigorous testing, should it be proven valid, translation into clinical trial has the potential to be rapid.

One question that requires consideration is what advantage does innervation bestow on the tumor? Nerves generally bundle along with blood vessels, providing ready access to required nutrients. Thus, it is possible that tumors induce their own innervation to provide a rich blood supply and promote tumor growth. Our data support this hypothesis. Consistent with this, EphrinB1 possesses proangiogenic properties (46) (47). Alternatively, tumor innervation may regulate the local immune response which in many solid cancers, particularly head and neck cancers, critically contributes to disease progression (48). Importantly, neuro-immune interactions are evolutionarily conserved and critical for homeostasis. Recent clinical trials using electrical stimulation of the vagus nerve demonstrate attenuation of disease severity in rheumatoid arthritis, an autoimmune disease (49). These and other data support the concept that alterations in neuroimmune interactions participate in disease pathogenesis and that therapeutic modulation of these interactions can restore homeostasis. Thus, tumors may promote their own innervation as a means to dampen immune responses, promote tumor tolerance, disease progression and dissemination. Examining the relationship between tumor innervation, vascularization and immune infiltrates in the tumor will help distinguish between these possibilities.

## Acknowledgments

This project was supported by an Institutional Development Award (IDeA) from the National Institute of General Medical Sciences of the National Institutes of Health under grant numbers 2P20GM103548 (Cancer), 5P20GM103620 (Pediatrics) and P20GM121341 (Population Health). Specifically, the Molecular Pathology Core, Imaging Core, Flow Cytometry and COMMAND (Collection Methods, Management and Analysis of Data) Cores provided their services and expertise towards this project.

The authors have no conflicts of interest to report.

Author contributions to this work are as follows:

M.M.: Experimental design, acquisition, analysis and interpretation of data, intellectual contributions and critical review of manuscript.

P.L.C.: Experimental conception and design, acquisition, analysis and interpretation of data, intellectual contributions and critical review of manuscript.

D.W.V.: Formulation of theory and prediction, experimental conception and design, acquisition, analysis and interpretation of data, intellectual contributions, critical review of manuscript.

C.T.L.: Acquisition, analysis and interpretation of data, critical review of manuscript.

E.G.V.: Acquisition, analysis and interpretation of data, critical review of manuscript.

A.J.G.: Acquisition and analysis of data.

J.T.C.: Experimental conception and design, acquisition and analysis of data.

D.M.: Analysis of data.

A.P.R.: Acquisition, analysis and interpretation of data.

Z.H.: Analysis and interpretation of data.

W.C.S.: Experimental conception and design, intellectual contributions, critical review of manuscript.

J.H.L.: Intellectual contributions, critical review of manuscript.

R.D.: Analysis and data interpretation, intellectual contributions, critical review of manuscript.

P.D.V.: Formulation of theory and prediction, intellectual contributions, experimental conception and design, drafting and revising of manuscript.

## Materials and Methods

### ANTIBODIES

<u>Antibodies utilized for western blot analysis included:</u> anti-CD9 (Abcam, 1:1,000), anti-CD81 (clone B-11, 1:1,000, Santa Cruz), anti-Ephrin-B1 (ECD epitope, R&D Systems, 1:500), anti-human EphrinB1 (ICD epitope, LifeSpan BioSciences, 1:500), anti-β-III Tubulin (2G10, 1:5,000, Abcam), anti-GAPDH (Ambion, 1:5,000). HRP- coupled secondary antibodies were purchased from Thermo-Fisher.

<u>Antibodies utilized for IHC:</u> anti-β-III Tubulin (2G10, 1:250, Abcam), anti-Tyrosine Hydroxylase (Ab112, 1:750, Abcam), anti-TRPV1 (cat# ACC-030, 1:100, Alomone labs), anti-VIP (ab22736, 1:100, Abcam).

<u>Antibody utilized for quantification on the CX7:</u> anti-β-III tubulin (Millipore, AB9354).

### CELL LINES

All cell lines have been authenticated by STR (BioSynthesis). In addition, all cell lines have been confirmed as mycoplasma free as per Uphoff and Drexler (50).

**<u>Human</u>**: UM-SCC1 and UM-SCC47 cell lines were maintained with DMEM with 10% fetal calf serum and 1% penicillin/streptomycin. Primary human tonsil epithelia were collected under an approved IRB protocol and maintained with KSFM (Gibco, cat # 10724-011). HTE E6/E7 and HTE E6Δ/E7 were generated by retroviral transduction and maintained in E-media as described above.

**<u>Mouse</u>**: mEERL cells (parental and all derivatives) were maintained with E-medium (DMEM (Corning, cat# 10-017-CV)/Hams F12 (Corning, cat#10-080-CV), 10% exosome depleted fetal calf serum, 1% penicillin/streptomycin, 0.5 µg/ml hydrocortisone, 8.4 ng/ml cholera toxin, 5 μg/ml transferrin, 5 µg/ml insulin, 1.36 ng/ml tri-iodo-thyonine, and 5 ng/ml EGF.

<u>mEERL EphrinB1 CRISPR clones:</u> Two distinct strategies were utilized to generate EphrinB1 null mEERL cell lines; one strategy employed simultaneous double-targeting to remove a large portion of gDNA spanning exons 1-5 (as in (*51*)) and one utilized single-targeting to produce frame-shift causing indels leading to early termination. Target selection and guide sequence cloning were carried out using the tools and protocol of Ran et al. (*52*). PCR assays for the double targeting strategy employed primers external to (1-5Δ Ext.) or within (1-5Δ Int.) the predicted deletion site (Table 1). The external assay should result in a 10,485bp wt amplicon and a 229bp Δ amplicon while the internal assay produces a 330bp wt amplicon and no Δ amplicon. Single target screening utilized PCR to amplify a 330bp region surrounding the target site followed by restriction digest with BslI, the recognition site of which should be destroyed when double strand breaks are incorrectly repaired (Table 1).

**TABLE 1.**
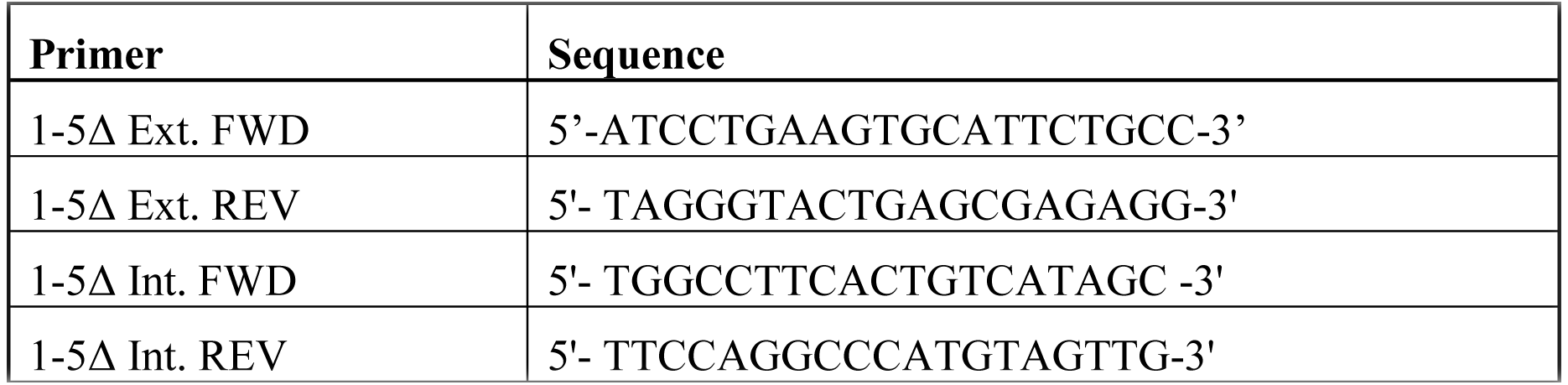
Primer sequences used to screen clones.

<u>mEERL EphrinB1ΔECD clone:</u> This clone was generated from the double-targeting strategy. PCR assays show the predicted deletion product using primers external to the targeted region and lack of an amplicon using primers within the deletion (Supplemental Figure 1B). The sequence data shows that Exons 2-4 are deleted; these exons comprise the majority of the extracellular domain of EphrinB1 (Supplemental Figure 1A). The 5’ end of the deletion in exon 1 occurs just after the signal peptide while the 3’ end of the deletion in exon 5 is within the transmembrane (TM) domain. Eight amino acids within the TM domain are deleted, however, two additional hydrophobic alanines are incorporated.

<u>mEERL Rab27 CRISPR clones:</u> Knockouts of Rab27 in mEERL cells were created using the general protocol of Ran et al. (52). Briefly, guide sequences targeting exons of RAB27A, RAB27B, or both were cloned into pSpCas9(BB)-2A-Zeo and transfected singly or in combinations to produce indels or larger deletions, respectively, in one or both genes. Following 5 days of Zeocin selection, single cells were expanded and screened for loss of restriction enzyme sites due to indels or by PCR for large deletions induced by double-targeting. Using this strategy, a single clone was identified for sequencing and further characterization.

<u>Clone mEERL Rab27A^−/+^Rab27B^−/−^:</u> This clone resulted from a strategy for double knock out of RAB27A and RAB27B in which sgRNA’s targeted to exon 4 of RAB27A, exon 4 of RAB27B, and a sequence of exon 3 shared by RAB27A and B were co-transfected. PCR revealed a heterozygous, truncated deletion product for Rab27A (Supplemental Figure 5A) and the expected homozygous deletion amplicon for Rab27B (Supplemental Figure 5B). RAB27B sequence data indicated distinct repair products at the deletion site; although one allele exhibits an immediate stop codon, it is unclear where the other might terminate. However, western blotting confirms lack of detectable protein (data not shown).

**<u>PC12 cells</u>**: PC12 cells were purchased from ATCC and maintained with DMEM with 10% horse serum (Gibco, cat # 26050-088) and 5% fetal calf serum. When used for neurite outgrowth assays, PC12 cells were maintained with DMEM with 1% horse serum and 0.5% fetal calf serum.

### IMAGING

<u>Electron microscopy:</u> Exosome samples were processed and analyzed by the Microscopy and Cell Analysis Core at Mayo Clinic: http://www.mayo.edu/research/core-resources/microscopy-cell-analysis-core/overview.

<u>Atomic Force Microscopy (AFM):</u> Purified exosomes were diluted 1:10 in de-ionized water, added to a clean glass dish, and allowed to air-dry for 2 hours before drying under a gentle stream of nitrogen. Exosomes deposited on glass dish were characterized using an AFM (Model: MFP-3D BIO^TM^, Asylum Research, Santa Barbara, CA). Images were acquired in AC mode in air using a silicon probe (AC240TS-R3, Asylum Research) with a typical resonance frequency of 70 kHz and spring constant of 2 Nm^−1^. Height and amplitude images were recorded simultaneously at 512 × 512 pixels with a scan rate of 0.6 Hz. Image processing was done using Igor Pro 6.34 (WaveMetrics, Portland, OR) and analyzed with Image J.

<u>Immunohistochemistry (IHC).</u> Tissues were fixed in 10% neutral buffered formalin and processed on a Leica 300 ASP tissue processor. All tissues were sectioned at 5 μm: N=12 human HNSCCs stained for TH, VIP and TRPV1 and N= 30 HNSCCs were stained for β-III tubulin. N=15 mEERL tumors were stained or TH, VIP and TRPV1 and N=30 mEERL tumors were stained for β-III tubulin. The BenchMark_®_ XT automated slide staining system (Ventana Medical Systems, Inc.) was used for the optimization and staining. The Ventana iView DAB detection kit was used as the chromogen and the slides were counterstained with hematoxylin. Omission of the primary antibody served as the negative control.

<u>PC12 assay and β-III tubulin quantification by CX7.</u> The CellInSight CX7 High Content Analysis Platform performs automated cellular imaging for quantitative microscopy which was utilized to quantify neurite outgrowth. 7.5×10^4^ PC12 cells were seeded onto 96 well black optical bottom, flat bottom plates (ThermoFisher) and 48 hours after treatment were fixed with 4% paraformaldehyde and then blocked and permeabilized with a solution containing 3% donkey serum, 1% BSA, and 0.5% Triton-X 100. Staining for β-III tubulin (Millipore, AB9354) was followed by Alexa Fluor^TM^ 488 goat anti-chicken IgG and Hoechst 33342. Washes were performed with PBS. Neurite outgrowth analysis was performed on the CellInsight CX7 HCS (ThermoFisher) using the Cellomics Scan Software’s (Version 6.6.0, ThermoFisher) Neuronal Profiling Bioapplication (Version 4.2). Twenty-five imaging fields were collected per well with a 10x objective with 2x2 binning. Nuclei were identified by Hoechst −positive staining, while cell somas and neurites were identified by β-III tubulin −positive immunolabeling. Cells were classified as neurons if they had both a Hoechst −positive nucleus as well as a β-III tubulin positive soma. Only neurites longer than 20 µm were included in the analysis. All assays utilizing exosomes from cell lines were run with an N=4 replicates per condition and repeated at least two times with similar results. Assays utilizing human samples were limited in materials and replicates were run to the extent possible as noted in the text.

### EXOSOME PURIFICATION

<u>Differential Ultracentrifugation.</u> 500,000 cells were seeded onto a 150 mm^2^ plate and incubated in medium containing 10% fetal calf serum that was depleted of exosomes. Fetal calf serum exosome depletion consisted of an over-night ultracentrifugation at 100,000 x g. Conditioned medium was collected after 48 hours and exosomes were purified by differential ultracentifugation as described by Kowal et al (31) with some modifications. Briefly, conditioned medium was centrifuged at 300 × g for 10 min at 4 °C to pellet cells. Supernatant was centrifuged at 2,000 × g for 20 min at 4 °C, transferred to new tubes, and centrifuged for 30 min at 10,000 × g, and finally in a SureSpin 630/17 rotor for 120 min at 100,000 × g. All pellets were washed in PBS and re-centrifuged at the same speed and re-suspended in 200 μL of sterile PBS/150mm dishes.

<u>Differential ultracentrifugation and optiprep density gradient.</u> Following differential ultracentrifugation as described above, a discontinuous iodixanol gradient was utilized similar to Van Deun et al (38) with some modifications. Solutions of 5, 10, 20 and 40% iodixanol were made by mixing appropriate amounts of a homogenization buffer [0.25 M sucrose, 1 mM EDTA, 10 mM Tris-HCL, (pH 7.4)] and an iodixanol solution. This solution was prepared by combining a stock solution of OptiPrep^TM^ (60% (w/v) aqueous iodixanol solution, Sigma) and a solution buffer [0.25 M sucrose, 6 mM EDTA, 60 mM Tris-HCl, (pH 7.4)]. The gradient was formed by layering 4 mL of 40%, 4 mL of 20%, 4 mL of 10% and 3 mL of 5% solutions on top of each other in a 15.5 mL open top polyallomer tube (Beckman Coulter). 400 µl of crude exosomes (isolated by differential ultracentrifugation) were overlaid onto the top of the gradient which was then centrifuged for 18 hours at 100,000 g and 4°C (SureSpin 630/17 rotor, ThermoScientific^TM^ Sorvall^TM^). Gradient fractions of 1 mL were collected from the top of the self-forming gradient, diluted to 14 mL in PBS and centrifuged for 3 hours at 100,000 g and 4°C. The resulting pellets were re-suspended in 100 µL PBS and stored at −80°C.

<u>Exosome purification from human plasma.</u> Ten ml of whole blood were pipetted directly onto Ficoll-loaded Leucosep tubes and centrifuged at room temperature for 30 minutes at 800 x g with the brake off. Exosomes were isolated from the recovered plasma by differential ultracentrifugation as described.

<u>Exosome purification from human tumor.</u> Fresh tumor tissue was cut into small pieces and placed in culture with KSFM (keratinocyte serum free medium) containing Fungizone (Thermo Fisher) and maintained in culture for 48 hours. Conditioned media was collected and exosomes harvested by differential ultracentrifugation as described.

### PROTEIN ANALYSIS

<u>BCA protein assay of exosomes.</u> The standard BCA protein assay was utilized with modifications to accommodate the low protein yield from exosome preparations. Briefly, 5μl of 10% TX-100 (Thermo Scientific) were added to an aliquot of 50 μl of purified exosomes and incubated 10 minutes at room temperature. A working ratio of 1:11 was used and incubated in a 96 well plate for 1 hr at 37°C. Absorbance at 562nm was then measured (SpectraMax Plus 384)and protein concentration estimated from a quartic model fit to the BSA standard curve.

<u>Western blot analysis.</u> Sample protein concentration was determined by BCA protein assay as described. Equal total protein was separated by SDS-PAGE, transferred to PVDF membranes (Immobilon-P, Millipore), blocked with either 5% Bovine Albumin Fraction V (Millipore) or 5% milk (Carnation instant non-fat dry milk), washed in TTBS (0.05% Tween-20, 1.37M NaCl, 27mM KCl, 25mM Tris Base), and incubated in primary antibody. Washed membranes were incubated with HRP-conjugated secondary antibody, incubated with chemiluminescent substrate (ThermoScientific, SuperSignal West Pico) and imaged using a UVP BioImaging System.

<u>***IN VIVO STUDIES.***</u> All animal studies were performed under approved institutional IACUC protocols and within institutional guidelines. All animal experiments utilized 4-8 week old male C57Bl/6 mice (The Jackson Laboratory) which were maintained at the Sanford Research Laboratory Animal Research Facility in accordance with USDA guidelines.

<u>Mouse tumor experiments:</u> Tumors were initiated as follows: using a 23-gauge needle, mEERL cells (1 × 10^5^ cells) were implanted subcutaneously in the right hind limb of mice. Tumor growth was monitored weekly by caliper measurements. Mice were euthanized when tumor volume was greater than 1.5 cm in any dimension. N=4 mice/group for quantification of β-III tubulin by western blot. N=7 mice/group for tumor growth.

<u>Whole tumor lysates.</u> Tumors were harvested 10 or 21 days post-implantation (as per text) and homogenized in lysis buffer on ice using a tissue homogenizer (Omni TH International). The homogenate was then sonicated and centrifuged at 2000 g for 5min. The resulting supernatant was collected and further centrifuged at 13000g for 10 min prior to BCA protein concentration estimation. Western blots were conducted using 30 µg inputs.

Beta-III tubulin western blots of whole tumor lysates from N=4 tumors/condition. Signals were quantified by densitometry using VisionWorks LS software and normalized to GAPDH. Group averages were compared using student’s t-test.

**<u>HUMAN SAMPLES.</u>** All human samples were collected under an approved Institutional Review Board protocol with signed Informed Consent. Samples included adult (age≥ 18 years) patients of both sexes and all races with a diagnosis of primary or locally advanced, squamous cell carcinoma of the head and neck (anatomic sites: oral cavity, oropharynx, hypopharynx, and larynx).

**<u>STATISTICAL ANALYSIS.</u>** Data were analyzed and graphed using PrismGraph. Descriptive statistics are presented as mean ± SEM or standard deviation (see Figure legends). Unpaired student’s t-test or one-way ANOVA were utilized for statistical analysis as indicated in the figure legends. PC12 assays utilizing exosomes from cell lines were run with four technical replicates for each condition and experiments were repeated at least 2 times. PC12 assays utilizing exosomes from human samples (blood or tumor) were treated differently as these samples were very limited. Thus, exosomes for each human sample were tested in duplicate when possible. When samples were limited (noted in text) only one well was tested.

